# A Meta-learning based Graph-Hierarchical Clustering Method for Single Cell RNA-Seq Data

**DOI:** 10.1101/2022.09.06.506784

**Authors:** Zixiang Pan, Yuefan Lin, Haokun Zhang, Yuansong Zeng, Weijiang Yu, Yuedong Yang

## Abstract

Single cell sequencing techniques enable researchers view complex bio-tissues from a more precise perspective to identify cell types. However, more and more recent works have been done to find more detailed subtypes within already known cell types. Here, we present MeHi-SCC, a method which utilized meta-learning protocol and brought in multi scRNA-seq datasets’ information in order to assist graph-based hierarchical sub-clustering process. In result, MeHi-SCC outperformed current-prevailing scRNA clustering methods and successfully identified cell subtypes in two large scale cell atlas. Our codes and datasets are available online at https://github.com/biomed-AI/MeHi-SCC

## I. Introduction

Single cell level sequencing was one of the most promising bio-technologies of this decade. It enables deep insight into cell-level biological process including cell communication and other micro-environment events[1, 2]. Nowadays, thanks to the progress of bio-engineering development and the next generation sequencing (NGS), single cell sequencing has already enabled us to detect signals via transcriptomes, DNA methylation and surface protein via micro-array chips[3, 4]. Among them, single cell RNA sequencing can be the most prevailing technology as it manifests more genetics information than other omics and have higher resolution compared to single cell DNA-seq with high performance-price ratio[5]. After fulfillment of Human Genome Project (HGP), larger scale cell atlas schedules were conducted like T cell atlas and B cell atlas, and quantity of cells were extended from 10^3^ to 10^6^[6–8]. However, technique limitations like sequence depth and systematic bias usually led to noise thus lower RNA counts sequencing accuracy like dropout[9]. At the same time, High dimension brought in challenges in dealing with batch effect and other differences between sequencing techniques, considering sometimes multi-source datasets are brought in for reference[10].

One primary task for single cell analysis is cell clustering. For now, single cell clustering helps detect new subtypes within known cell types, which helps reveal bio-mechanism in oncology micro-environment heterogeneity and tissue cell differentiation in embryo genesis[11, 12]. To tackle problems mentioned above, dimension reduction methods like LSI were developed[13], and low-dimension visualization methods like T-SNE, and UMAP are helpful for providing researchers a direct version into these high-redundant and high-dimensional datasets[14, 15].

Representation learning, aroused by dimension reduction, is now an important realm in analyzing single cell data in artificial intelligence, especially in coping with problems like imputation, cell type annotation and clustering[16–19]. Eraslan et al. first applied auto-encoder to extract deep low-dimensional space for single cells with Zero-inflated Negative Binomial distribution[9], while Tian et al. first used learnt latent space to generate clustering labels for further self-supervision with auto-encoder[20]. Xie et al. utilized dual self-supervised techniques to assign high dimension samples with several clustering centers[21], to better preserve local data feature structure, Guo et al. proposed IDEC, which preserved auto-encoder structure in dual self-supervision training phase[22]. To better reveal inter-cellular relationship, Zeng et al. integrated cell-cell relationship with graph neuron network as extra information to optimize dual self-supervised clustering[23]. In addition, Yi et al. applied graph attention network enhancement for the same goal and Hui et al. brought in contrastive learning to promote robustness[24, 25]. As for traditional machine learning based single cell clustering method, Seurat uses log-normalization to impute raw RNA counts and generate clustering labels based on nearest neighbors with high variant gene markers[26]. Wang et al. applied multi-kernel learning for better cell representation learning[27], while Lin et al. promoted principal component analysis and imputation, and is scalable to cluster thousands of cells[28].

However, traditional self-supervised clustering methods have limitations considering data sparsity and dataset specificity. One way to cope with the problem is to bring in more bio-medical information or multi-datasets as reference, and meta-learning is a brand new field to do so. Meta-Learning, namely “learning to learn”, is a brand new concept based on large data and multi-task training, which was considered as a substitution with traditional fine-tuning, and now it has gained preference in many tasks in computer vision and nature language processing realms[29–31].

In this paper, we proposed a Meta-learning based Hierarchical Single Cell Clustering tool called MeHi-SCC. The main motivation of MeHi-SCC is to explore a potential way to cluster a certain scRNA dataset into more detailed sub-types with the aid of hierarchical clustering. Inspired by Xing et al.[32], we used meta-training and brought in multi clustering tasks with multi-source scRNA datasets and their former manually annotated cell types as support tasks to promote clustering accuracy.

MeHi-SCC have three main innovating contributions as follows:

1. MeHi-SCC features meta-learning. We trained an independent hierarchical clustering separator section LANDER with support clustering tasks[32]. Unseen scRNA datasets from different sequencing techniques, tissues or even different species were fed to LANDER one by one for training with annotated ground truth. Noticeably, support datasets have no overlap with query dataset.
2. MeHi-SCC features a whole-graph-tuning based hierarchical clustering section. LANDER, the separator, only learns how inter-cellular relationship helps cluster step by step toward ground truth, ignoring specific expression values. Different from graph convolution neural-network(GNN) with fixed adjacent matrix, LANDER updates both edge-connections and related node features.
3. MeHi-SCC enables sub-cell-type detection. Hierarchical LANDER helps divide cell graphs into sub-cell graphs and aggregate them into more detailed clusters for all cells until they cannot be divided into sub-graphs any more, and our cluster number is usually more than ground truth given by manual annotations from morphology or biomarkers.

In this work, we proved that MeHi-SCC performed better than current-prevailing benchmark pipelines. Besides, MeHi-SCC sees better latent representation visualization performance compared with other pipelines and we successfully identified cell sub-types in two large scale single cell atlas, from both human and mouse tissue.

## II. Materials and Methods

MeHi-SCC mainly separates into three parts(Fig.1), i.e. de-noising auto-encoder part, meta-learning part, and dual sself-supervision part. All the parts are based on Pytorch framework[33]. De-noising auto-encoder part, as shown on the left-top of the figure, used for data pre-processing for both support and query datasets, is actually a set of auto-encoders with Gaussian noise added. Meta-training part, as shown on the right-top of the figure, is hierarchical LANDER, a meta-learning based hierarchical clustering separator machine.

**Fig.1.**
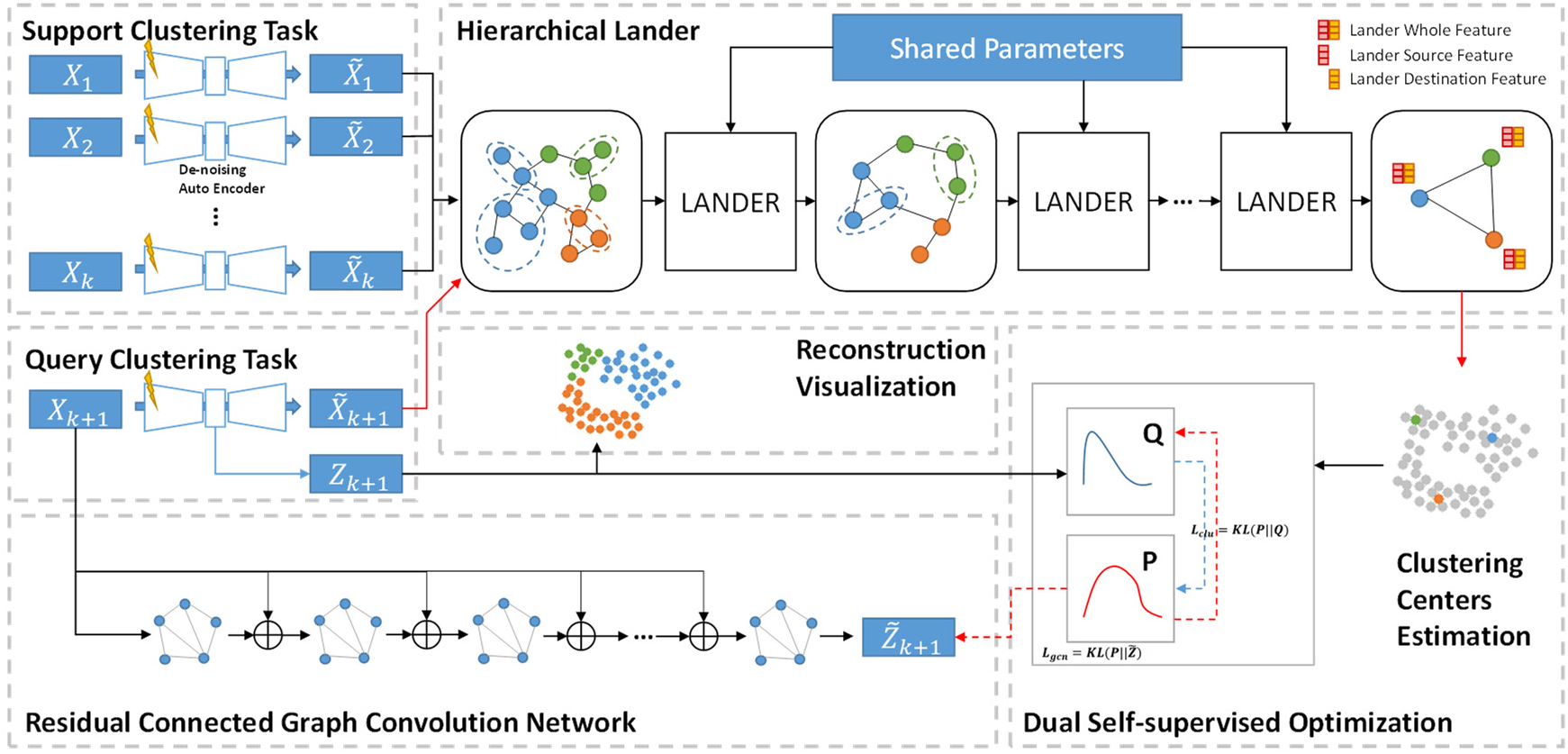
Basic Framework of MeHi-SCC.

Datasets should first be reconstructed by the de-noising auto-encoder. And then reconstructed expression profiles will be fed to well pre-trained LANDER as input to generate initial clustering results(red arrows).

Finally the dual self-supervision part, on the bottom of the figure, will calculate clustering centers from clusters generated by LANDER, and then latent spaces generated by the de-noising auto-encoder and the residual connected graph convolution network (Left Down) will approach to the distribution of clustering centers through Kullback-Leibler Divergence[34].

### A. Data Preparation and De-Noising Auto-Encoder

We here prepared 15 open-sourced datasets for experiments, with 13 normal scale datasets and 2 large-scale datasets(PBMC 68K and Shekhar Mouse Retina). All the datasets have their manual annotated cell types, we downloaded their expression profiles and treated the cell type annotations as ground truth(Supplementary Materials Table S1).

Before scRNA-seq profiles were fed to the model, we first applied raw counts (including the query dataset and support training datasets) with TPM normalization via Seurat[26]. And then we selected 2000 High Variant Genes (HVGs) by default. Then the 2000-dimension-profile will be fed to the de-noising auto-encoder by 400 epochs and then to the LANDER.

Here, the origin scRNA profile *X* ∈ ℝ^*N*×*m*^ has *N* cells and *m* selected genes, and Gaussian noise *Δ* has the same dimension with *X*. The auto-encoder *h* consists of encoder *h*_*e*_ with dimensions [*m*, 512,256,64, *z*] where *z* denotes latent dimension. *Z* = *h*_*e*_(*X* + *Δ*) and *Z* ∈ ℝ^*N*×*z*^ is the latent space representation which is used for visualization, and then the latent space will be used as input of decoder *h_d_* with dimensions [*z*, 64,256,512, *m*] to calculate reconstruction expression profile 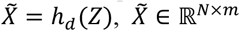. The optimization goal for de-noising auto-encoder is mean square error(MSE), i.e. 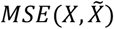.

### B. Training the Meta Learning LANDER

LANDER is a Graph Convolution Network (GNN) based framework with two part —— the separator *ϕ* and aggregator *ψ*. The separator *ϕ* consists of learnable a Graph Attention Network (GAT), which is used for separating graph into sub-graphs[35], and aggregator *ψ* play the role as aggregating sub-graphs to clustering labels according to sub-graphs.

The meta-training phase of LANDER was conducted on multi clustering tasks one by one. For example, we have *n*_*train*_ training tasks, and each task has a single cell transcriptional expression profile with manually annotated ground truth. At first, after pre-processed and de-noised, we obtained *i*^*th*^ reconstructed expression profile 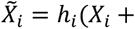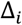) with Gaussian noise *Δ*_*i*_. The *i*^*th*^ auto-encoder *h*_*i*_ consists of encoder and decoder mentioned above, and then we built K-nearest neighbor (KNN) with DGL toolkit to build graph for every reconstructed expression 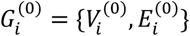 where origin nodes 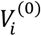 are cells whose node features 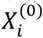 are reconstructed expression, i.e. 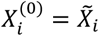, and edges 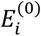 are determined by k nearest neighbors[36]. Noticeably, LANDER is dismountable and was pre-trained independently from rest parts of MeHi-SCC framework.

As a hierarchical clustering method, multi shared-parameters LANDERs are applied to divide current KNN graphs into sub-graphs layer by layer. Here we define 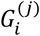 as the graph of the *i*^*th*^ training task/loop and the output of *j*^*th*^ LANDER. The *j*^*th*^ LANDER (also called “ *j*^*th*^ level”) has its separator *ϕ*^(*j*)^. Edges are subset of the input graph edges indicating that 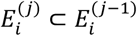 and are calculated via the separator that 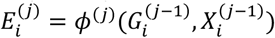. The new graph will be constructed based on split edge components 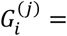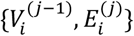. Then aggregator *ψ* will be utilized to calculate node features of the next level, i.e. 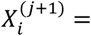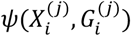. Here the aggregator compress every connected nodes pair into a single node for the next level. The construction of new edge set 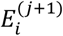 is still based on KNN searching on 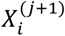. When no more edges are added, i.e. 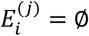, the split reaches convergence (i.e. early stopping). Finally LANDER will conclude and assign every cell with a clustering label according to which sub-graph it belongs to from the input level to the convergence level.

To be more precise, LANDER conducts clustering job without reliance on specific features, but on relationship between features and their neighborhood edges, i.e. the graph. Thus graph densities and linkages matter. For each edge we calculate density via two vertices (*v*_*m*_, *v*_*n*_) ∈ *E* and linkage probabilities *p*_*mn*_ = *P*(*y*_*m*_ = *y*_*n*_) by a Multi Layer Perceptron (MLP) network with Softmax as the output transferring function. Their concatenated feature [*x*_*m*_, *x*_*n*_] of the vertices pair (*v*_*m*_, *v*_*n*_) is used as input of the MLP network. Here *y*_*m*_ is the manually annotated cell type labels of *v*_*m*_ given from the support datasets. The node density 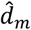 of cell *v*_*m*_ was calculated within *k* nearest neighbors for every node as 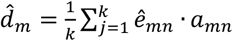, where 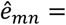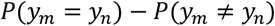 is the edge coefficient and *a*_*mn*_ is the inner product of node features between vertices pair (*v*_*m*_, *v*_*n*_), which is defined as *a*_*mn*_ = 〈*x*_*m*_, *x*_*n*_〉

Separator *ϕ* is used to update edge connections for LANDER output. With pre-given connection latch *p*_*τ*_ (tuned during meta-training phase and stay fixed after meta-training, and for test clustering label estimation, the parameter is usually set 0.1), potential cell set for *v*_*m*_ is defined as 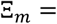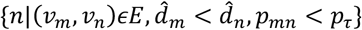, and final edges linked with the cells that 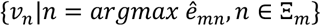 are under the precondition that *Ξ*_*m*_ ≠ ∅.

After the *j*^*th*^ separator *ϕ*^(*j*)^ separates a certain graph *G*^(*j*)^ into *P* sub-parts 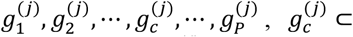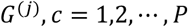, the *c*^*th*^ sub-part 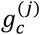 will be first converted to a single node 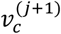 for the next level 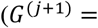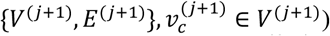, whose feature consists of two concatenated parts 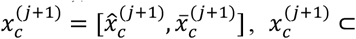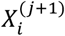. Here 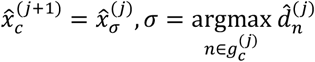 is the source feature who represents the peak node feature (the node with highest density) from the related sub-graphs, and 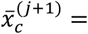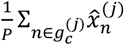 is the destination feature which denotes average feature of the related sub-graph 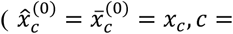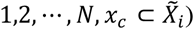.

Here, suppose *d*_*m*_ as cell density calculated by ground truth cell type label and ground-truth-based edge coefficients are calculate by *e*_*mn*_ = *I*(*y*_*m*_ = *y*_*n*_) − *I*(*y*_*m*_ ≠ *y*_*n*_) where *I*(·) is the indicator function. The two optimization goals in the meta-training phase are on nodes and edges respectively. The first is the density loss function defined as below to approximate estimated cell densities to ground truth densities:

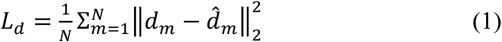

Another optimization goal has restriction on edges, i.e. average edge cross-entropy to form nodes on each side of an edge linkage to a same cluster:

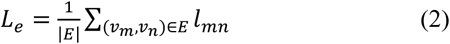

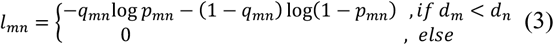

Then the total loss of LANDER meta training is:

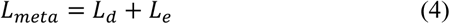

### C. Dual Self-Supervision for Further Clustering

Finally, after LANDERs are pre-trained with meta-training phase, we then apply residual graph convolution to further optimize clustering results. We constructed graph from dot-product similarity matrix *S* ∈ ℝ^*N*×*N*^ with de-noising encoder embedding *Z*_*AE*_ = *h*_*e*_(*X*), *Z*_*AE*_ = {*z*_1_, *z*_2_, …, *z*_*m*_, …, *z*_*n*_, …, *z*_*N*_}^*T*^, where *z*_*m*_ ∈ ℝ^*z*^, *m* = 1,2, …, *N*. We calculate similarity between two cells, 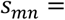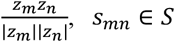:

Here we use similarity matrix to select *k*′ (*k*′ = min{0.01*N*, 20}) cells as neighbors to construct graph adjacent matrix *A*. The residual graph is introduced in to make up for structure information that the de-noising auto-encoder missed:

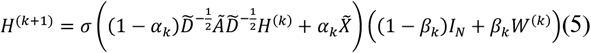

Where *σ* depicts activation function RELU, parameters are *α*_*k*_ = 0.3 and *β*_*k*_ = 0.5 × (*k*′)^−1^ according to reference[37]. Normalized degree matrix 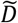 is calculated via adjacent matrix with self-loop 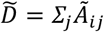 and 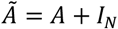.

We then obtained final latent space of the residual graph network as a confidence matrix *Z*_*g*_ = *soft*max(*W*^(*k*)^*H*^(*k*)^ + *b*^(*k*)^), *Z*_*g*_ ∈ ℝ^*N*×*c*^ and *c* is the number of cell types given by hierarchical LANDER.

From pre-proposed research, Student’s t-distribution helps optimize clustering. We fetched clustering labels given by LANDER as *Y*_*O*_ and calculated the *l*^*th*^ clustering centers *μ*_*l*_ for each cluster as 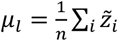.

Where 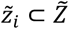 is latent cell space whose clustering label is *y*_*l*_ and *y*_*l*_ ∈ *Y*_*0*_. Hereby we are able to calculate similarity between the *m*^*th*^ cell and *l*^*th*^ clustering center as *q*_*ml*_, which can be regarded as the confidence degree (or the possibility) that the *m*^*th*^ cell belongs to the *l*^*th*^ cluster:

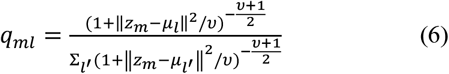

Here *υ* denotes the freedom degree and is set 1 by default. And then the assistant distribution is introduced in for promotion:

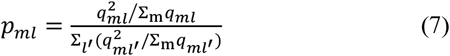

In all, supposed *P* = [*p*_*ml*_] and, the gold of minimizing the difference between the soft distribution *P* and the confidence degree matrix *Q* via KL divergence (Formula (8)) helps separation of representative learning. The similarity between soft distribution *P* to graph-network-learnt embedding *Z* acts likewise (Formula (9)). Thus the final training loss function is be defined as (10):

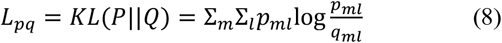

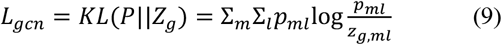

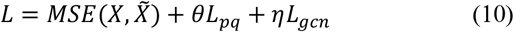

Here, *z*_*g,ml*_ ⊂ *Z*_*g*_, and *θ* is set as 0.1 and *η* as 0.01 for all query test datasets. After 1000 epochs of joint training with (10), the final general clustering result *Y*′ of cell *m* will be calculated as 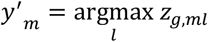.

Noticeably, *Y*′ shows further optimized aggregated clustering labels from LANDER clusters, while *Y*″ generated from formula (7) manifests more detailed sub-clusters within *Y*′. *Y*″, whose components *y*″_*m*_ are calculated as follows (*p*_*g,ml*_ have similar definition with *z*_*g,ml*_ and are components of *P*), is regarded as detailed sub-cluster result as 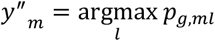.

## III. Results

### A. Performance on 14-1 Meta Training

Here we evaluate performance between benchmark methods on 13 normal scale datasets, and MeHi-SCC shows great superiority. Before dual self-supervision module, we trained LANDER for every query task with other 14 datasets’ clustering tasks as support tasks within all our 15 datasets. The meta-training phase bases on multi-Ks (i.e.[4, 5, 38]) for DGL graph construction and multi-levels (i.e.[1, 3, 4]). The meta-training’s learning rate is set 0.01 with stochastic gradient descent optimizer and batch size is set 4096 for 1000 epochs for each support tasks[39].

After LANDER’s training was done, we applied several Ks (ranging from 3 to 10) and levels (ranging from 2 to 5) to test their potential highest performance. In terms of the dual self-supervision module, we first applied LANDER on reconstructed expression profiles after 400 epochs of de-noising pre-training to fetch the LANDER clustering labels and related clustering centers. The 400 epochs pre-training was based on Adam optimizer[40].

We mainly use three indices for evaluation, i.e. Adjusted Rand Index (ARI), Normalized Mutual Information (NMI) and Clustering Accuracy (CA), calculated by sklearn package[41] (Supplementary Materials Formula S1-S3) .

In order to prove the effectiveness of meta-learning, we removed meta-training phase and only preserved hierarchical clustering in our framework with other conditions fixed (“MeHi-SCC without meta-learning” in Table I.), and the performance sees significant decrease without meta-learning.

**TABLE I.** Average Performance Comparison on 13 Normal Scale Datasets

| Methods | ARI | NMI | CA |
| --- | --- | --- | --- |
| CIDR[28] | 0.4955 | 0.6476 | 0.6195 |
| DCA[9] | 0.5035 | 0.6306 | 0.5989 |
| IDEC[22] | 0.4708 | 0.5888 | 0.5952 |
| scDeepCluster[20] | 0.5600 | 0.7299 | 0.6433 |
| Seurat[26] | 0.5792 | 0.7474 | 0.6765 |
| SIMLR[27] | 0.5368 | 0.6721 | 0.6518 |
| scGAC[24] | 0.6319 | 0.7268 | 0.7277 |
| scNAME[25] | 0.6447 | 0.7192 | 0.7476 |
| MeHi-SCC without meta-learning | 0.6031 | 0.6820 | 0.7148 |
| **MeHi-SCC** | **0.7698** | **0.7752** | **0.8119** |

Generally, MeHi-SCC outperformed all its counterpart methods(Table I.). In Fig.2, we plot all ARI, NMI and CA performance with violin plots. On each plot, each violin bar includes 13 points indicating 13 normal scale datasets. Finally, MeHi-SCC obtained 0.7698 ARI on average (Fig.2(a)), exceeding scNAME (0.6447) by 19.4%, and scGAC (0.6319) by 21.8%. The average NMI were 0.7752 (Fig.2(b)), outperformed Seurat (0.7474) by 3.7% and scDeepCluster (0.7299) by 6.2%. The average CA finally reached 0.8119 on average(Fig.2(c)), which is 8.6% higher than scNAME (0.7476) and 11.6 % higher than scGAC (0.7277).

**Fig. 2.**
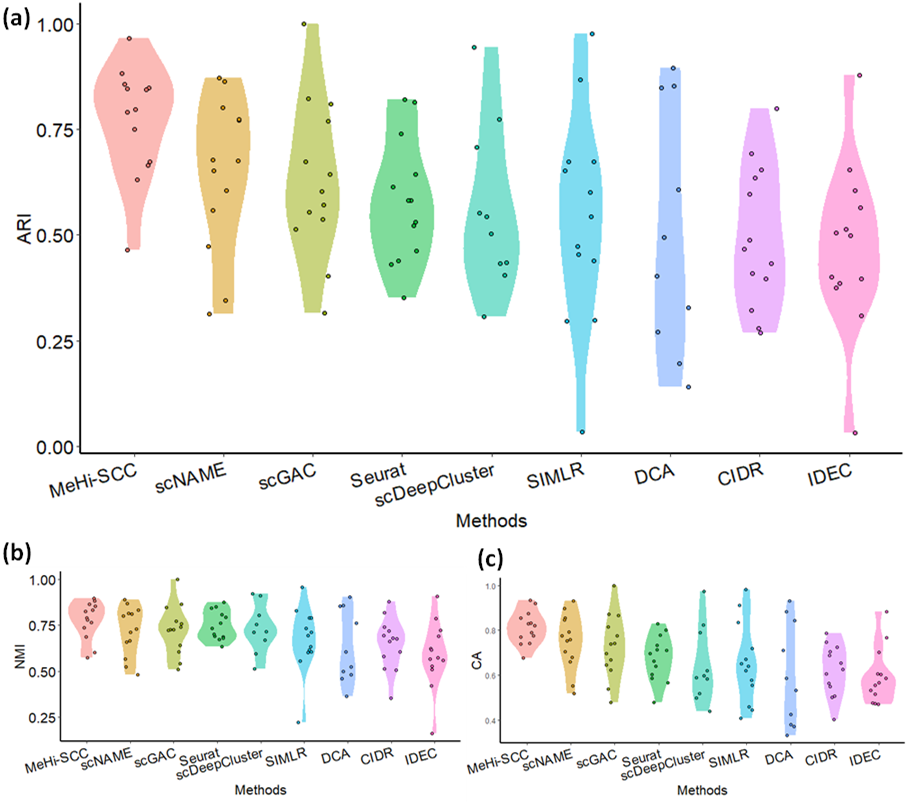
Overall Performance on 13 Normal Scale Datasets. (a) ARI, (b) NMI, (c) CA.

### B. Ablation Experiment

The connection between LANDER and the following dual self-supervision module plays an important role in overall performance. Thus we examined influence of different kinds of clustering centers towards overall performance. The five types of clustering centers includes:

1. LANDER Whole Feature + PCA: We reduced dimension of the last layer (level) of LANDER graph features to the same dimension as the latent auto-encoder representations with principle component analysis (PCA)[42].
2. LANDER Src./Dst. + PCA: Since the source feature and the destination feature, are the same in the last level, we fetched them for PCA dimension reduction. Different from LANDER Whole Feature, the source/destination feature only has half of the LANDER whole feature dimension.
3. LANDER Whole Feature + AE. Here we reduced LANDER whole feature dimension with the de-noising auto-encoder we pre-trained instead of PCA to the latent shape as clustering centers.
4. LANDER Clusters Average. This usage is what MeHi-SCC takes. Different from other types of centers mentioned about, here we do not use LANDER features as clustering centers but only use LANDER clustering results whose geometric centers are considered as clustering centers.
5. K-means. The clustering centers of K-means are actually geometric centers of each clusters calculated by K-means. For fair comparison, we set K in K-means the same to the number of clusters calculated by LANDER.

All the ablation experiments were conveyed on same conditions, and then we measured average performance for 13 normal scale datasets (Table II). From UMAP visualization (Supplementary Materials Fig.S1), we spot that graph features generated by LANDER, with lower variance and regardless of dimension reduction methods, hardly fit in with other cells so that they cannot be used as clustering centers(red spots). In contrast, both K-means centers and LANDER clusters average centers performed better in that their clustering centers evenly allocated inside the entire cell manifolds. Thanks to the merged information from meta-learning, LANDER clusters average centers exceeded K-means by 15.3%, 5.5% and 11.4% on ARI, NMI and CA respectively.

**TABLE II.** Ablation Experiments on Different Clustering Center Types

| Clustering Center Type | ARI | NMI | CA |
| --- | --- | --- | --- |
| LANDER Whole Feature + PCA | 0.3412 | 0.5610 | 0.4533 |
| LANDER Src./Dst. Feature + PCA | 0.2266 | 0.4503 | 0.3568 |
| LANDER Whole Feature + AE | 0.3320 | 0.5459 | 0.4448 |
| **LANDER Clusters Average** | **0.7815** | **0.7827** | **0.8150** |
| K-means | 0.6776 | 0.7419 | 0.7316 |

### C. Visualization for Clustering Latent Spaces

Then we fetched embedding representation of three representative benchmark pipelines, scNAME, scGAC and Seurat, together with MeHi-SCC for visualization with ground truth. Latent space from MeHi-SCC learnt a good representation with clear separation between basic cell types (Fig.3). We retrieved latent embedding spaces from MeHi-SCC de-noising auto-encoder after the whole training phase. For the rest of the benchmarks, we utilized default parameters. All the methods was visualized by UMAP. For those benchmarks based on R, we applied package umap to calculate dimension reduced coordinates, and for benchmarks based on python sessions, we visualized their latent space via packages scanpy and anndata[43].

**Fig. 3.**
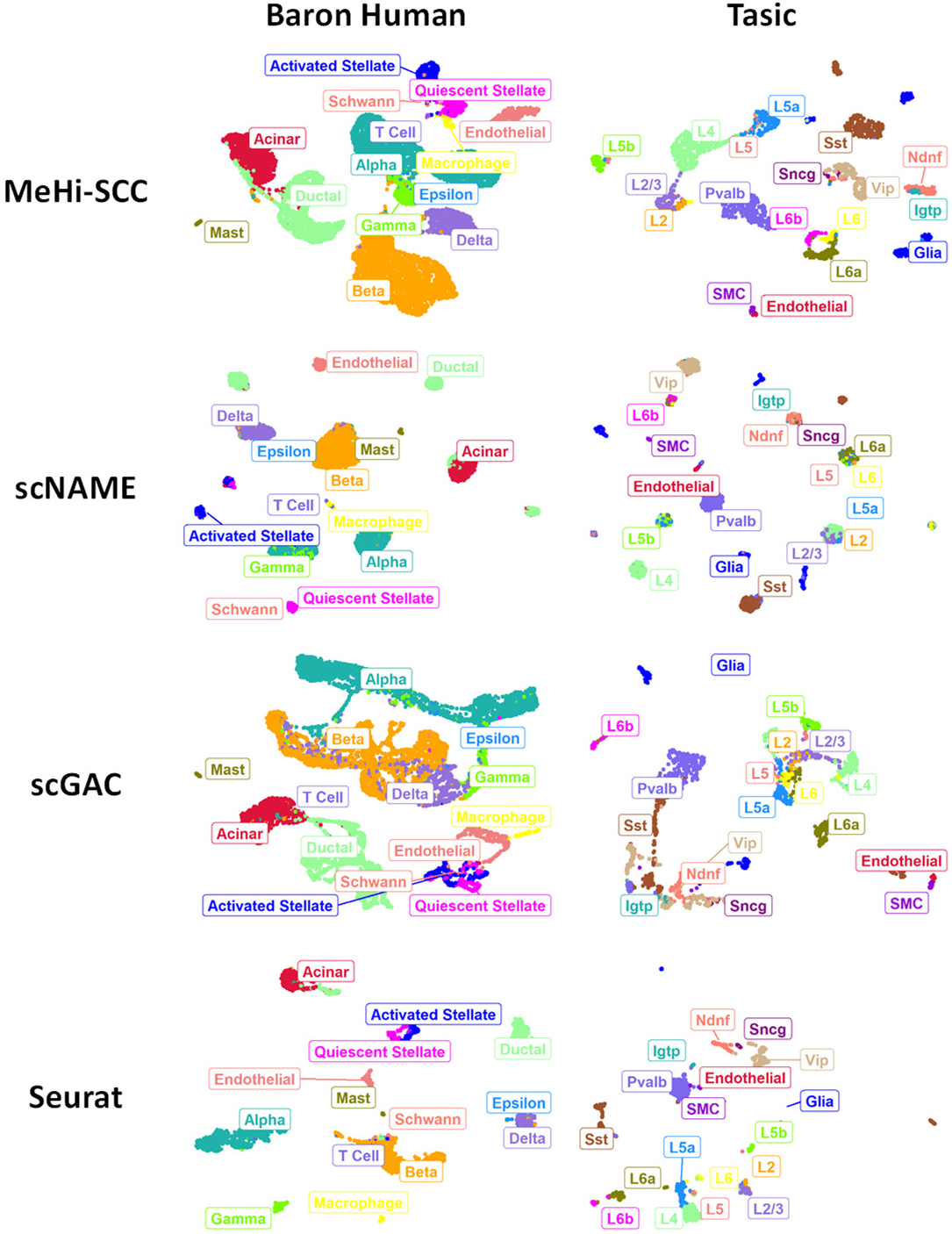
Separation Performance Comparison on Latent Space of MeHi-SCC and Three Representative Methods on two example datasets with UMAP.

MeHi-SCC sees clear separation on Baron Human dataset, compared with other benchmark methods. In scNAME latent space, Beta cells are mixed with mast cells, and activated stellate cells are mixed with quiescent stellate cells. The later phenomenon is also shown in scGAC latent space. In terms of Seurat, it generally performed well except that T cells are blended with Beta cells. Additionally, when it comes to Tasic, a small scale dataset, most of the benchmark pipelines separate cells well, except that, for scGAC, it is difficult to separate L2, L2/3, L5 and L5a cells.

### D. Case Studies of Sub-Cell Types Detection

Since MeHi-SCC is a hierarchical method, here we give two case studies of successful subtype detection. We applied gene enrichment analysis based on Seurat v3[26], then we retrieved detailed clustering results generated by MeHi-SCC on two large scale dataset PBMC 68K and Shekhar Mouse Retina.

After log-normalization and 2000 high variable genes selected, we then scaled the pre-normalized expression profiles, and ran PCA analysis to find neighbors with top 10 sPCs. Then we ran FindAllMarkers function in Seurat with min.pct=0.25 and filtered genes with log fold change threshold >1 and P value under 0.01 in each cluster. We here visualized the two datasets with latent spaces from MeHi-SCC, with ground truth labels (Fig.4(a), Supplementary Materials Fig.S2(a)) and detailed MeHi-SCC clustering results(Fig.4(b), Supplementary Materials Fig.S2(b)). After low quality clusters were filtered, we then referred to CellMarker database to validate potential clusters as cell sub-types[44].

In PBMC 68K, We found that CD37 enriched in cluster 10 and cluster 1[45], and MS4A1 enriched in cluster 10 indicating that cluster 10 could be identified as memory B cells among the CD19+ mature B cells[46]. GZMK is enriched in cluster 2 and cluster 23 indicating that we can identified effector memory CD8+ T cell inside CD8+ cytotoxic T cells[47]. Additionally, we found MZB1 and TNFRSF17 are enriched in cluster 20, seeing that MeHi-SCC deciphered the isolated part of the CD19+ B cells as dividing plasma B cells (Fig. 4(c))[45, 48, 49]. In term of Shekhar Mouse Retina, we found that Nfia is highly expressed especially in cluster 28 which is a potential mark for Rheaume et, al. 28 cells. Likewise, Gjc1 enriched in cluster 27 depicting markers of Rheaume et, al. 11 cells(Supplementary Materials Fig.S2(c))[50].

**Fig. 4.**
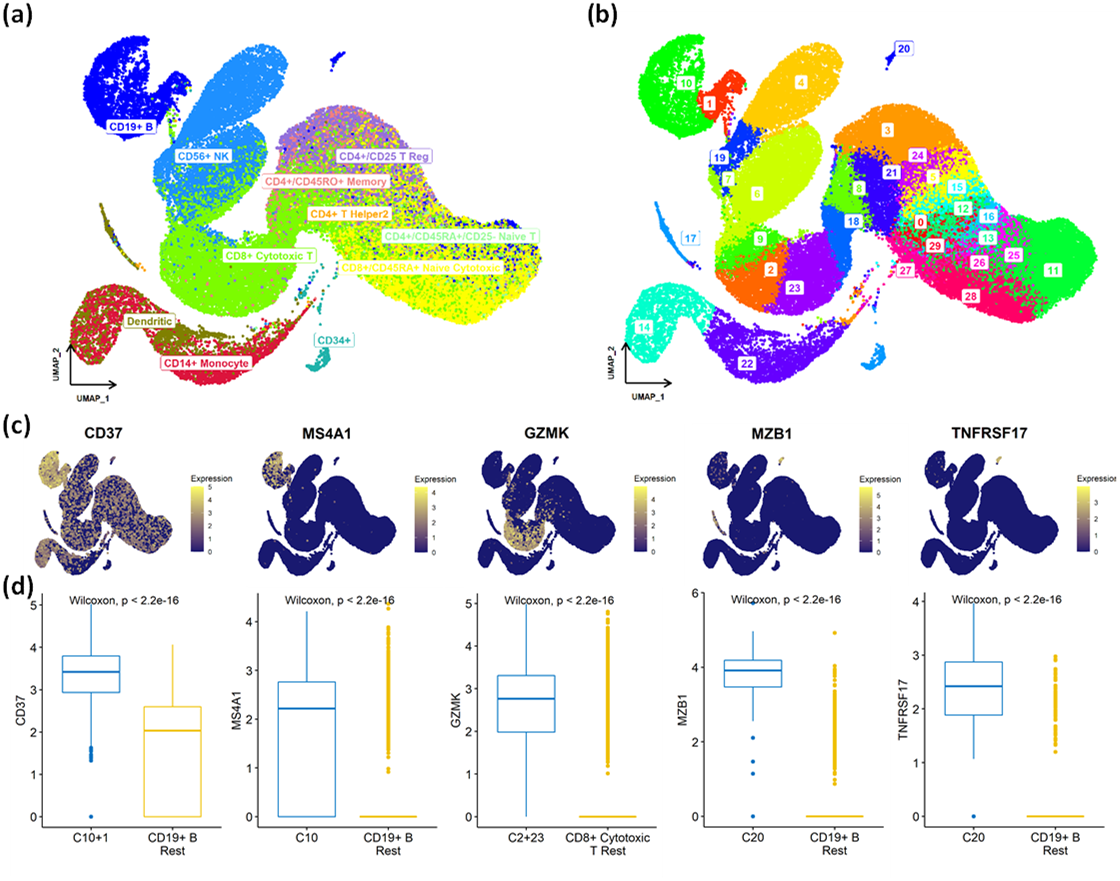
Cell Subtype Detection on PBMC 68K by MeHi-SCC. (a) Ground Truth Cell Types of PBMC 68K, (b) Clustering Results Given by MeHi-SCC on PBMC 68K, (c) Five Potential Marker Genes Visualized on PBMC 68K, (d) Significance Tests for CD37, MS4A1, GZMK, MZB1 and TNFRSF17.

We then ran Wilcoxon signed rank test on marker genes detected between MeHi-SCC clusters mentioned above and the rest part of cells who belong to the same ground truth cell types with the clusters above. We spot that all potential genes expressed are significantly expressed, with P-value under 0.01(Fig.4(d), Supplementary Materials Fig.S2(d)).

## IV. Discussion and conclusion

In summary, we proposed MeHi-SCC, a meta-learning based graph hierarchical clustering framework for scRNA-seq data. In this work, we bring in extra information from multi-source scRNA datasets to aid clustering. MeHi-SCC outperformed current prevailing single cell clustering methods and the hierarchical clustering mechanism helps decipher sub-cell-types within known cell types in large-scale cell atlas.

Additionally, we encourage following researchers to test whether meta-learning clustering can be applied to more sparse single cell omics data like scATAC data. With more information provided by whole genomic sequencing, scRNA-seq could be an ideal assistant in referring clusters for other omics. And whether we can use multi bulk transcriptional profiles as assistance for single cell analysis with aid of meta-learning deserves exploring.

## Acknowledgment

This work was sponsored by National Natural Science Foundation of China (61772566) and Guangdong Key Field R&D Plan (2019B020228001).

## Supplementary Information

Codes, example data and supplementary materials are available at https://github.com/biomed-AI/MeHi-SCC.

